## Supplementary Information for "Actin Dysregulation Induces Neuroendocrine Plasticity and Immune Evasion: A Vulnerability of Small Cell Lung Cancer"

### Supplementary Figures

Supplementary Figure 1. CRACD inactivation in SCLC
Supplementary Figure 2. scRNA-seq of preSC (*Cracd* WT vs. KO) allograft tumors
Supplementary Figure 3. scRNA-seq of RPR2 vs. CRPR2 SCLC tumors
Supplementary Figure 4. NOTCH signaling downregulation by *Cracd* KO
Supplementary Figure 5. Immune cell profiling of RPR2 vs. CRPR2 SCLC tumors
Supplementary Figure 6. scRNA-seq-based immune cell profiling
Supplementary Figure 7. Impact of EZH2 blockade on Cracd-inactivated SCLC tumorigenesis
Supplementary Figure 8. scRNA-seq analysis of the human SCLC tumor datasets

**Supplementary Video 1**. Prediction of cell lineage trajectories of murine SCLC tumors (RPR2 vs. CRPR2; preSC *Cracd* WT vs. KO allografts)

### Uncropped immunoblot images

### Supplementary Tables

Supplementary Table 1. Reagent information
Supplementary Table 2. Primers used for qRT-PCR
Supplementary Table 3. Top 30 genes of each cluster in RPR2 and CRPR2 scRNA-seq dataset
Supplementary Table 4. Top 30 genes of each cluster in preSC WT and preSC Cracd KO scRNA-seq dataset
Supplementary Table 5. Software and algorithms used for scRNA-seq analyses
Supplementary Table 6. Gene list used for EZH2 and PRC2 target scoring
Supplementary Table 7. Clinical information of human SCLC samples analyzed by scRNA-seq
Supplementary Table 8. Information of human normal lung samples for scRNA-seq

### Supplementary Figures

#### Supplementary Figure 1. CRACD inactivation in SCLC

- **a.** Somatic alterations in *CRACD*, *TP53*, *RB1*, *RBL2*, *CREBBP*, and *EP300* in 249 SCLC patient samples. The diagram was generated using OncoPrinter at www.cbioportal.org/oncoprinter.jsp).
- **b.** Types of *CRACD* mutations found in SCLC patients and cell lines.
- **c.** Distribution and frequency of *CRACD* mutations found in 102 SCLC cell lines and 249 SCLC patient samples, illustrated along the length of protein. The diagram was generated using the Broad Institute Cancer Cell Line Encyclopedia [CCLE] and cBioPortal databases.
- **d.** Levels of *CRACD* mRNA transcripts in normal lung and SCLC patient tumor samples from the TCGA database; Student's *t*-test.

#### Supplementary Figure 2. scRNA-seq of preSC (*Cracd* WT vs. KO) allograft tumors

- **a.** UMAP of integrated scRNA-seq datasets of allograft tumor generated from *Cracd* WT vs. KO preSCs.
- **b.** UMAPs of subsets of cells from the global level (left) to the allograft tumor cells (right). Each dot represents a single cell, colored by cell type.
- **c.** Feature plots of mouse cell-type marker gene expression: *Epcam* (epithelial cells), *Ptprc* (immune cells), *Pecam1* (endothelial cells), and *Colla1* (mesenchymal cells).
- **d.** Heatmap of gene expression-based cell clusters. The top 10 genes that were highly expressed in each cell cluster were visualized; these were used for cell annotation of *Cracd* WT and KO allograft tumors.

#### Supplementary Figure 3. scRNA-seq of RPR2 vs. CRPR2 SCLC tumors

- **a.** UMAPs of the scRNA-seq datasets of normal mouse lung, RPR2, and CRPR2 SCLC tumors.
- **b.** UMAPs of the scRNA-seq datasets of RPR2 and CRPR2 SCLC tumors.
- **c.** UMAPs of subsets of cells from the global level (left) to SCLC tumor epithelial cells (right). Each dot represents a single cell, colored by cell type. Tumor epithelial cells (RPR2 and CRPR2) were replotted into another UMAP (right).
- **d.** Feature plots displaying the expression of each representative marker: *Epcam* (epithelial cells), *Ptprc* (immune cells), *Pecam1* (endothelial cells), *Colla1* (mesenchymal cells), and *Fn1* (mesenchymal cells).
- **e, f.** A copy number variation analysis of normal mouse lung, RPR2, and CRPR2 SCLC tumors. Copy number variation plot showing the distribution of genomic alterations (gains and loss) in RPR2, and CRPR2 SCLC tumors compared with healthy mouse lung samples (E). Copy number variation scores were projected into the UMAP of the scRNA-seq dataset from healthy human lung, RPR2, and CRPR2 datasets (F).
- **g.** Dot plots for mouse lung epithelial marker gene expression in each cell cluster.
- **h.** Heatmap of each cell cluster of RPR2 and CRPR2 tumors. The top 10 genes that were highly expressed in each cell cluster were visualized; these were used for cell annotation.
- **i.** Comparison of proportions of different cell types between the RPR2 and CRPR2 datasets.
- **j.** RNA velocity–based cell lineage trajectory analysis of RPR2 and CRPR2 scRNA-seq datasets. RNA velocity was calculated through a dynamic model, and cells were clustered using the "Leiden" algorithm, the scVelo and Scanpy packages (n_neighbors = 10, n_pcs = 40).
- **k.** Partition-based graph abstraction–based visualization of cell lineage trajectories of RPR2 and CRPR2 tumor cells. The size of the circle corresponds to the cell number. A partition-based graph abstraction analysis was performed and plotted using the RNA velocity–based cell clusters (scVelo). While clusters 4 and 10 in RPR2 tumors were inferred as root cell clusters, CRPR2 tumors likely harbored three additional root cell clusters (1, 2, and 9), repopulating tumor cells (also see Supplementary Video S1). Among those, cluster 1 was exclusively found in *Cracd* KO tumors (CRPR2). It should be noted that cell lineage trajectories were inferred by RNA splicing (RNA velocity) of scRNA-seq datasets.
- **l.** PHATE mapping of RPR2 and CRPR2 scRNA-seq datasets.

#### Supplementary Figure 4. NOTCH signaling downregulation by *Cracd* KO

**a.** IHC of murine lungs (RPR2 vs. CRPR2) for HES1.

**b.** IHC of murine lungs isolated from LUAD models (*Kras*$^{LSLG12D}$, *Trp53*$^{floxed/floxed}$ [KP]) instilled with adenovirus encoding Cre recombinase, Cas9, and sgRNAs (*LacZ* [control] or *Cracd*), showing the impact of conditional KO of *Cracd* on KP-driven LUAD tumorigenesis, as we recently performed[1].

**c.** IHC of murine lungs isolated from LUAD models (*Kras*$^{LSLG12D}$, *Trp53*$^{floxed/floxed}$ [KP] or *Cracd*$^{-/-}$, *Kras*$^{LSLG12D}$, *Trp53*$^{floxed/floxed}$ [CKP]) instilled with adenovirus encoding Cre recombinase, displaying the impact of germline KO of *Cracd* on KP-driven LUAD tumorigenesis, as we recently performed[1].

**d.** IB of RPR2 or CRPR2 cells treated with DAPT, a gamma-secretase inhibitor (10 uM, 48hrs). FL: full length, TM: transmembrane.

**Supplementary Figure 5. Immune cell profiling of RPR2 vs. CRPR2 SCLC tumors**

**a.** UMAPs of cells from the global level (left) and immune cell subsets (right) in RPR2 and CRPR2 scRNA-seq datasets. Each dot represents a single cell, colored by cell type.

**b.** UMAP of re-clustered whole immune cells of integrated RPR2 and CRPR2 datasets (left). Each cluster represents different immune cell types. Feature plots displaying the expression of T cell marker *Cd3d* and *Cd3g*; B cell marker *Cd79a;* NK cell marker Nkg7; Myeloid cell marker *Lyz2* and *Cd68* (right).

**c.** Dot plots show mouse immune cell marker gene expression in each cell cluster of integrated scRNA-seq datasets of *Cracd* WT and KO SCLC tumors.

**d.** Heatmap of gene expression–based cell clusters of the top 10 genes in each immune cell type of integrated RPR2 and CRPR2 datasets.

**e.** Analysis of tumor-infiltrated CD8$^+$ T cells. Immunostaining of RPR2 and CRPR2 tumors with anti-CD8 antibody (left). Nuclei were counterstained with DAPI. Quantification of CD8$^+$ T cell counts per 660 $\mu m^2$ in RPR2 and CRPR2 tumor tissue (right). Representative images are shown. *P* values were calculated using the Student's *t*-test; error bars: SD.

**f-h**. RPR2 and CRPR2 tumor tissues were stained with CD3, a marker for T cells, and cleaved caspase-3, a marker for cell apoptosis (F), and counted (G, H) per 660 $\mu m^2$ in RPR2 and CRPR2 tumor tissues. Nuclei were stained with DAPI. Representative images are shown. *P* values were calculated using the Student's *t*-test; error bars: SD. *Cracd* KO does not significantly affect the total T cell number or cell death.

**i, j**. No impact of *Cracd* loss on PD-1-PD-L1/2 signaling. Feature plots of *Pd-l1*, *Pd-l2*, and *Pd-1* expression in all cells between RPR2 and CRPR2 scRNA-seq datasets (I). Feature plots of *Pd-l1*, *Pd-l2*, and *Pd-1* expression in immune cells between RPR2 and CRPR2 datasets (J).

**k**. Dot plot showing the expression level of MDSC marker genes (*Itgam*, *Cd14*, *Clec4d/e*, *Il1b*, *Arg2*, *Wfdc17*, and *Cd300ld*) in RPR2 and CRPR2 datasets.

**l**. Upregulation of MDSC marker expression in myeloid cell cluster by *Cracd* KO. Feature plots of MDSC marker gene expression in immune cells between RPR2 and CRPR2 datasets. *Cracd* KO upregulates the expression of MDSC markers (*Itgam*, *Cd14*, *Clec4d/e*, *Il1b*, *Arg2*, *Wfdc17*, and *Cd300ld*). Circle, myeloid cell cluster.

**Supplementary Figure 6. scRNA-seq-based immune cell profiling**

**a.** UMAP of subsets of cells from the global level (upper) and immune cell subsets (lower) in integrated scRNA-seq datasets of preSC allograft tumors (*Cracd* WT vs. KO). Each dot represents a single cell, colored by cell type.

**b.** Heatmap of gene expression–based cell clusters of the top 10 genes that were most highly expressed in each immune cell type of integrated scRNA-seq datasets of preSC allograft tumors (*Cracd* WT vs. *Cracd* KO).

**c.** Dot plots show mouse immune cell marker gene expression in each cell cluster.

**d.** *Cracd* KO decreases the number of Cd8$^+$ effector T cells in preSC tumors. UMAPs of preSC *Cracd* WT (left) and preSC *Cracd* KO (right) subsets showing each immune cell type.

**e.** Cell proportion analysis of different immune cell types between preSC *Cracd* WT and KO datasets.

**f.** Dot plot showing the upregulation of MDSC markers expression by *Cracd* KO preSC tumors compared to *Cracd* WT preSC tumors. Circle, myeloid cell cluster.

**g.** Total cell-cell interactions (left) and interaction strength (right) from RPR2 and CRPR2 tumors were analyzed using the CellChat package.

**h.** Cell-cell interaction analysis with CellChat. Chord plots show significant changes in signaling between immune cells in the RPR2 (left) and CRPR2 (right) SCLC tumors. The inner bar colors represent the cell clusters that receive signals. The inner bar size is proportional to the signal strength received by the cell clusters. Chords indicate ligand-receptor pairs that mediate the interaction between two cell clusters; the size of the chords is proportional to the signal strength of the given ligand-receptor pair. RPR2 tumors showed a strong interaction between the MHC-I pathway (*H2-K*, *H2-D*, *H2-Q*, and *H2-T* from all tumor cell clusters) and CD8 receptors (from $CD8^+$ T cells). In contrast, CRPR2 tumors exhibited no interaction between the MHC-I pathway and CD8 receptors. Intriguingly, in CRPR2 tumors, significant cell-cell interactions include tumor cells (via App, Mif, and Ptn)-B cells (*Cd74_Cxcr4*), Naïve T cells (*Ncl*, *Cd47*), NK cells (*Ncl*, *Itga4*, *Cd44*, and *Cd47*), and macrophages (*Cd74*, *Cd44*, *Ncl*, and *Itga4*).

**Supplementary Figure 7. Impact of EZH2 blockade on SCLC tumorigenesis**

**a.** Cell viability assay of RPR2 and CRPR2 cell lines treated with EZH2 inhibitors, GSK343 and Tazemetostat. Cell viability was measured by Cell Counting Kit-8.

**b.** Flow cytometry analysis using FlowJo to assess T cell populations in CRPR2 tumors treated with vehicle (Veh), Tazemetostat (Taze), or GSK343 (GSK). Representative plots show pacific blue-CD4 or APC-CD8 T cell populations within PE-CD45 and FITC-CD3 double-positive cells. Q2 and Q6 quadrants indicate the percentage of positive cells.

**c.** Tumor growth curves of RPR2 subcutaneous tumors in immunocompetent mice treated with vehicle (Veh), Tazemetostat (Taze; 200 mg/kg, oral gavage), or GSK343 (GSK; 20mg/kg, intraperitoneal), administered every other day (n=10 per group). Tumor volume was measured every other day.

**d.** Tumor growth was subsequently assessed by measuring tumor weight. No statistical significance was observed; NS, not significant by Student's *t*-test ($P \geq 0.05$).

**Supplementary Figure 8. scRNA-seq analysis of the human SCLC tumor datasets**

**a, b.** A copy number variation analysis of MS1 and MS2 tumors. Copy number variation plot showing the distribution of genomic alterations (gains and loss) in MS1 and MS2 tumors compared with healthy human lung samples (A). Copy number variation scores were projected into the UMAP of the scRNA-seq dataset from healthy human lung samples and MS1/2 samples (B).

**Supplementary Video 1. Prediction of cell lineage trajectories of murine SCLC tumors (RPR2 vs. CRPR2; preSC *Cracd* WT vs. KO allografts)**

Animation of the reconstructed predicted cell fates and the most probable path of cell-state transitions shown in Figure 3 (RPR2 vs. CRPR2; upper panels and preSC *Cracd* WT vs. KO; lower panels). The Dynamo package was used to analyze gene expression dynamics and cell-state transition data. The red digit is emitting fixed points (initial cell states, stem cells); the blue digit is unstable fixed points (bifurcation, where the cell makes fate decisions, progenitor cells); the black digit is absorbing fixed points (terminal cell state, differentiated cells); the full circle is red or black fixed points, a half circle is blue fixed points; filled color of each fixed point represents the confidence, following the value of viridis color map. Yellower, higher value, more confident; greener, lower value, less confident.

### Uncropped immunoblot images

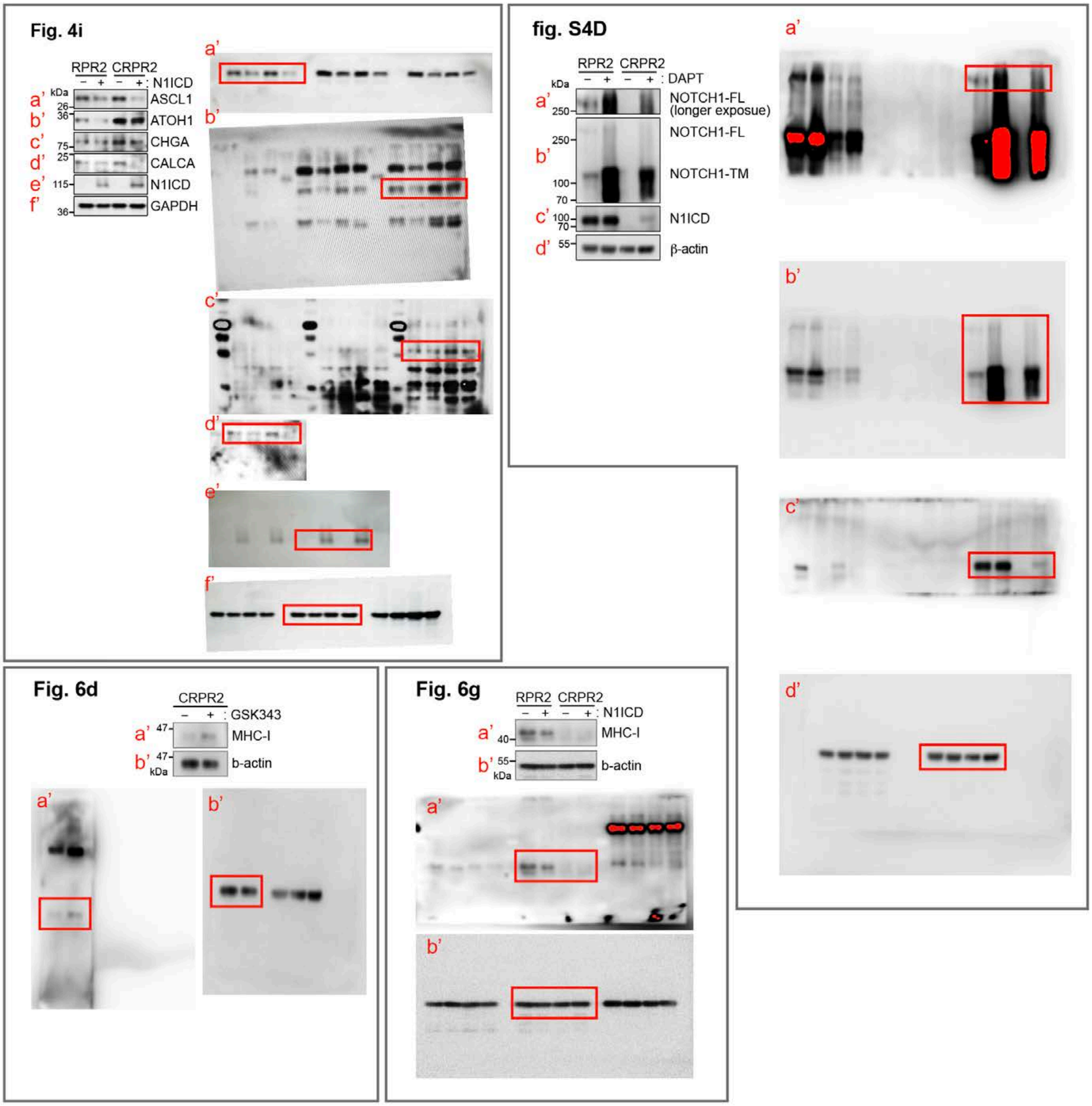


### Supplementary references

1. Kim B, Zhang S, Huang Y, Ko KP, Jung YS, Jang J, Zou G, Zhang J, Jun S, Kim KB, Park KS, Park JI. CRACD loss induces neuroendocrine cell plasticity of lung adenocarcinoma. Cell Rep. 2024;43(6):114286. Epub 20240525. doi: 10.1016/j.celrep.2024.114286. PubMed PMID: 38796854; PMCID: PMC11216895.
