## Supplementary Figures for "Actin Dysregulation Induces Neuroendocrine Plasticity and Immune Evasion: A Vulnerability of Small Cell Lung Cancer"

**Supplementary Figure 1**

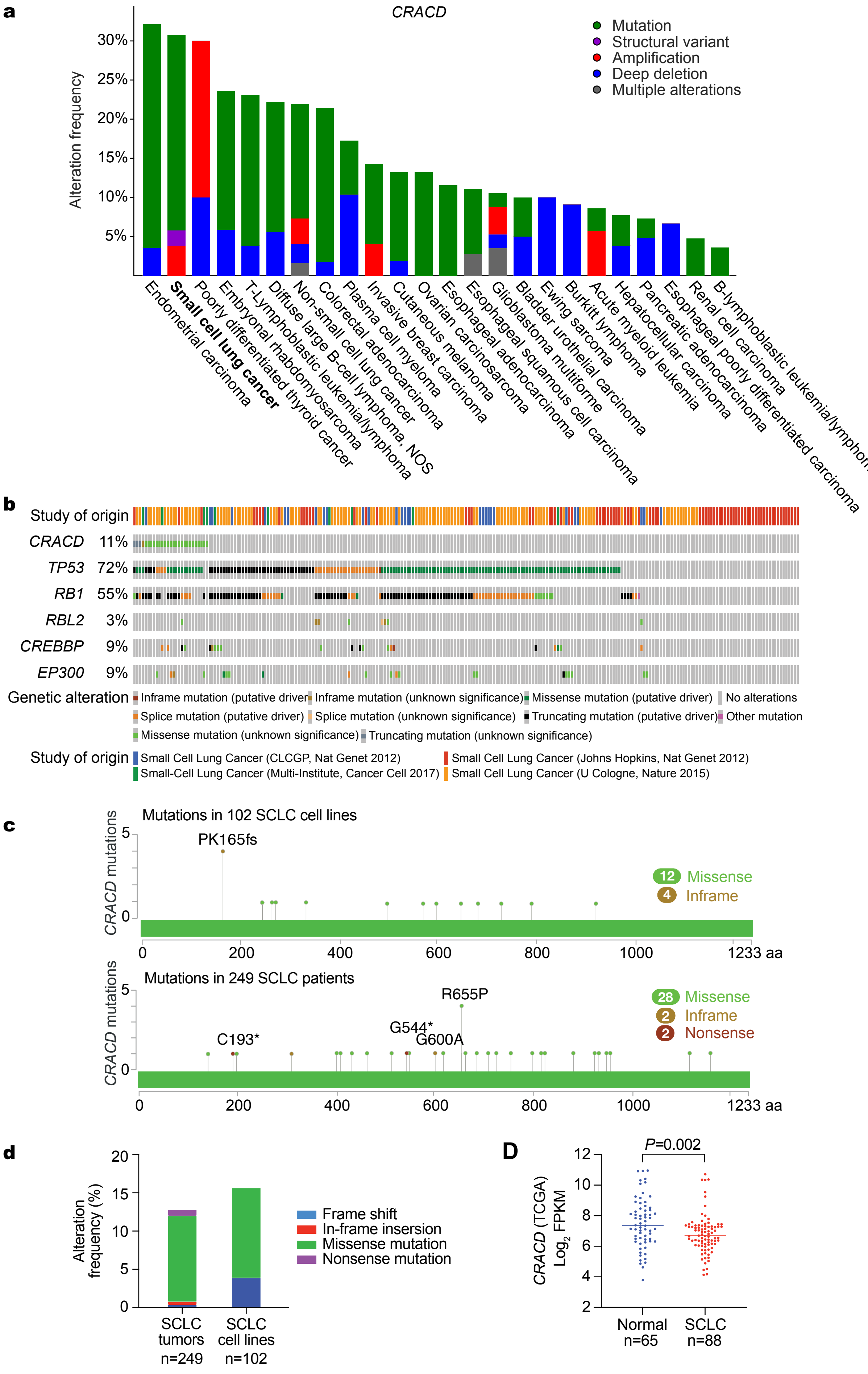

### Supplementary Figure 2

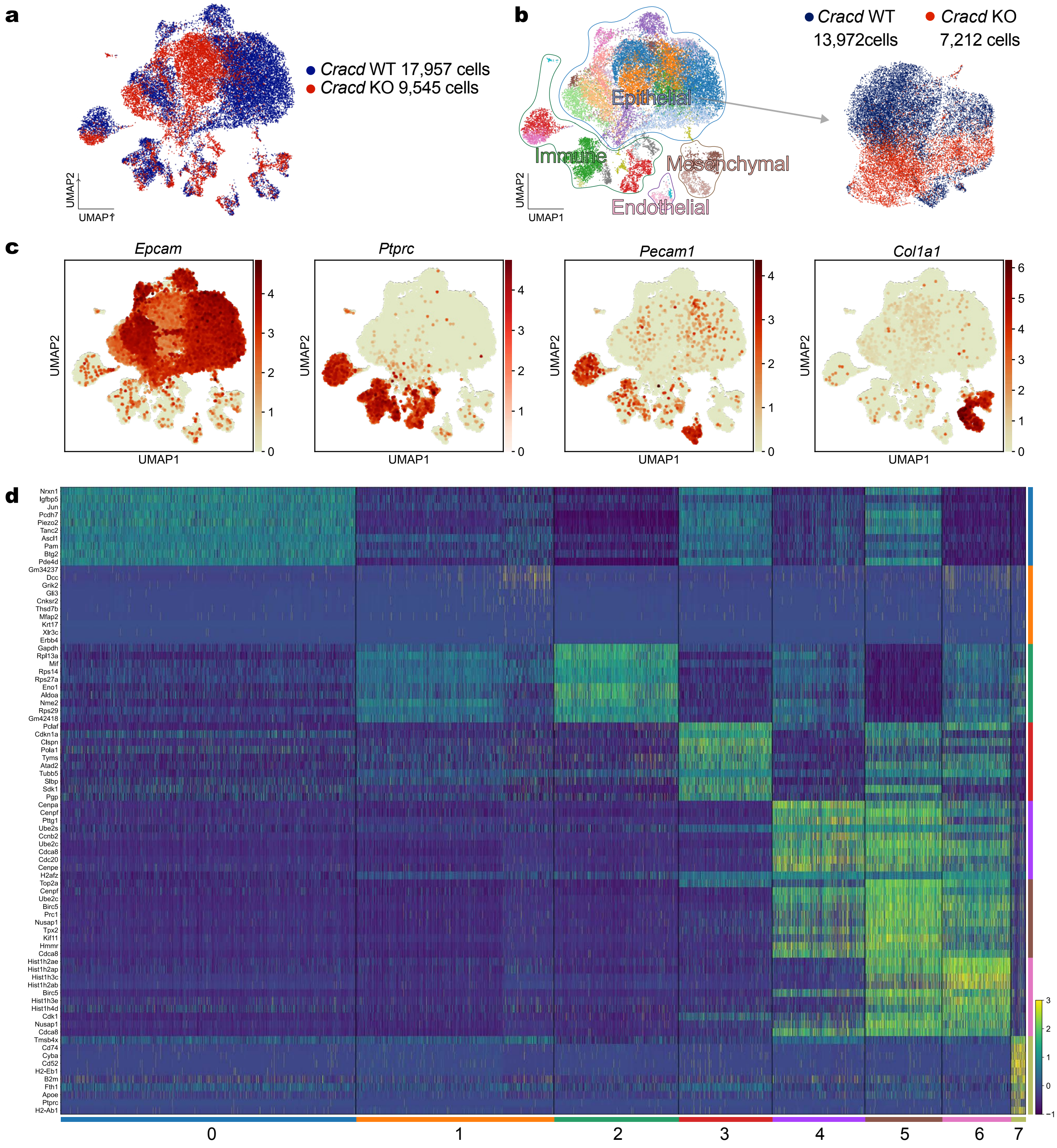

**Supplementary Figure 3**

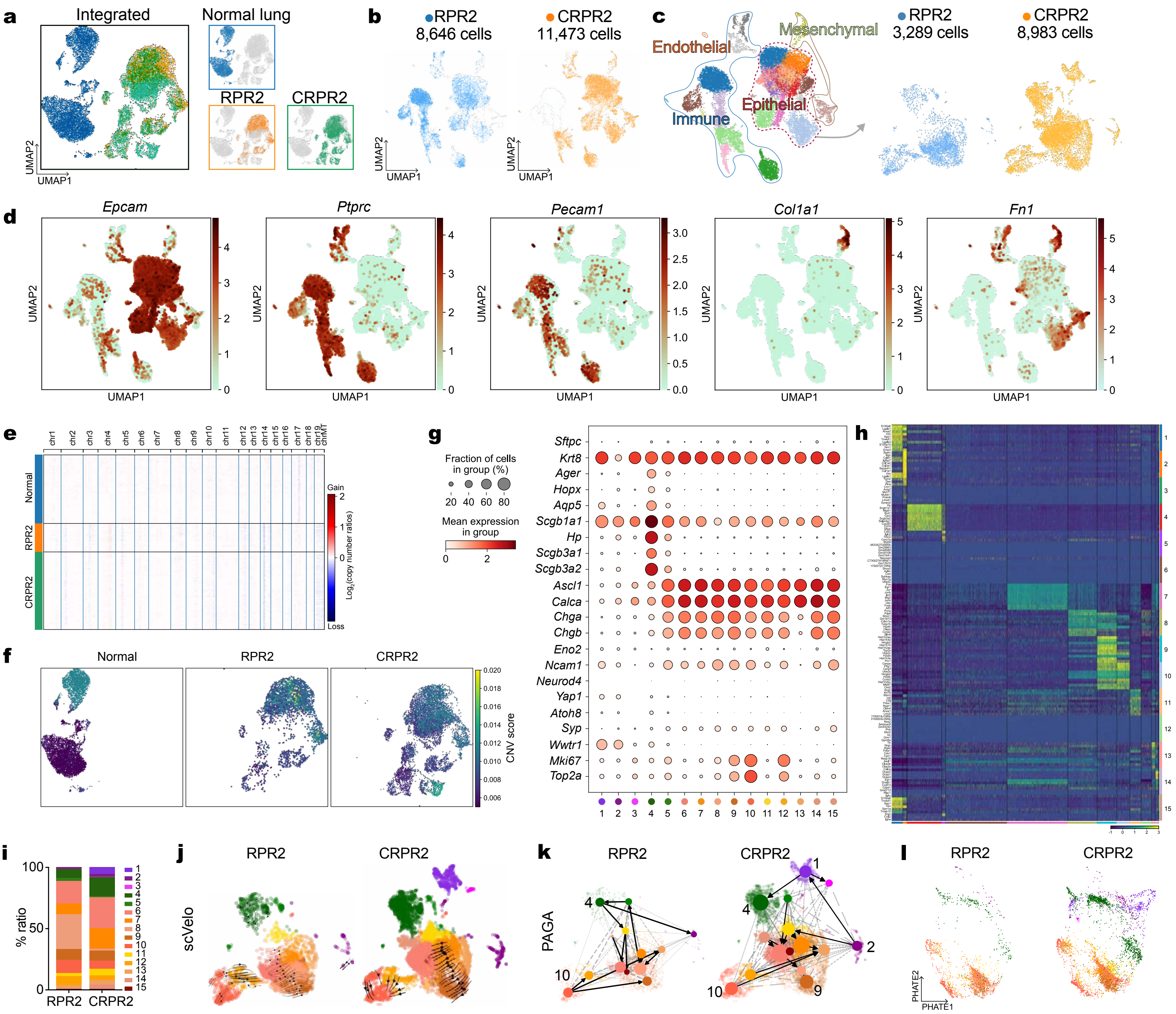

**Supplementary Figure 4**

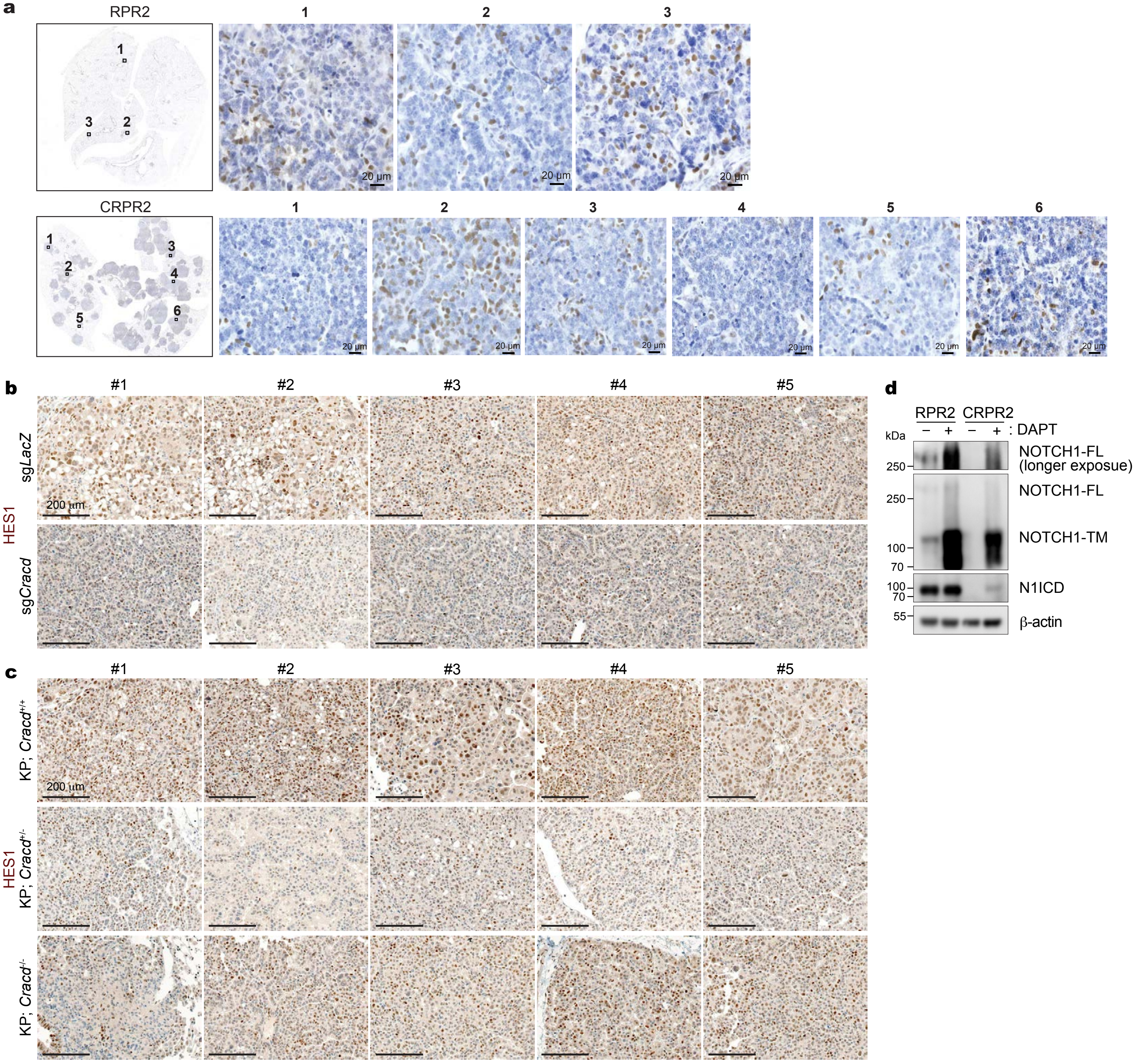

**Supplementary Figure 5**

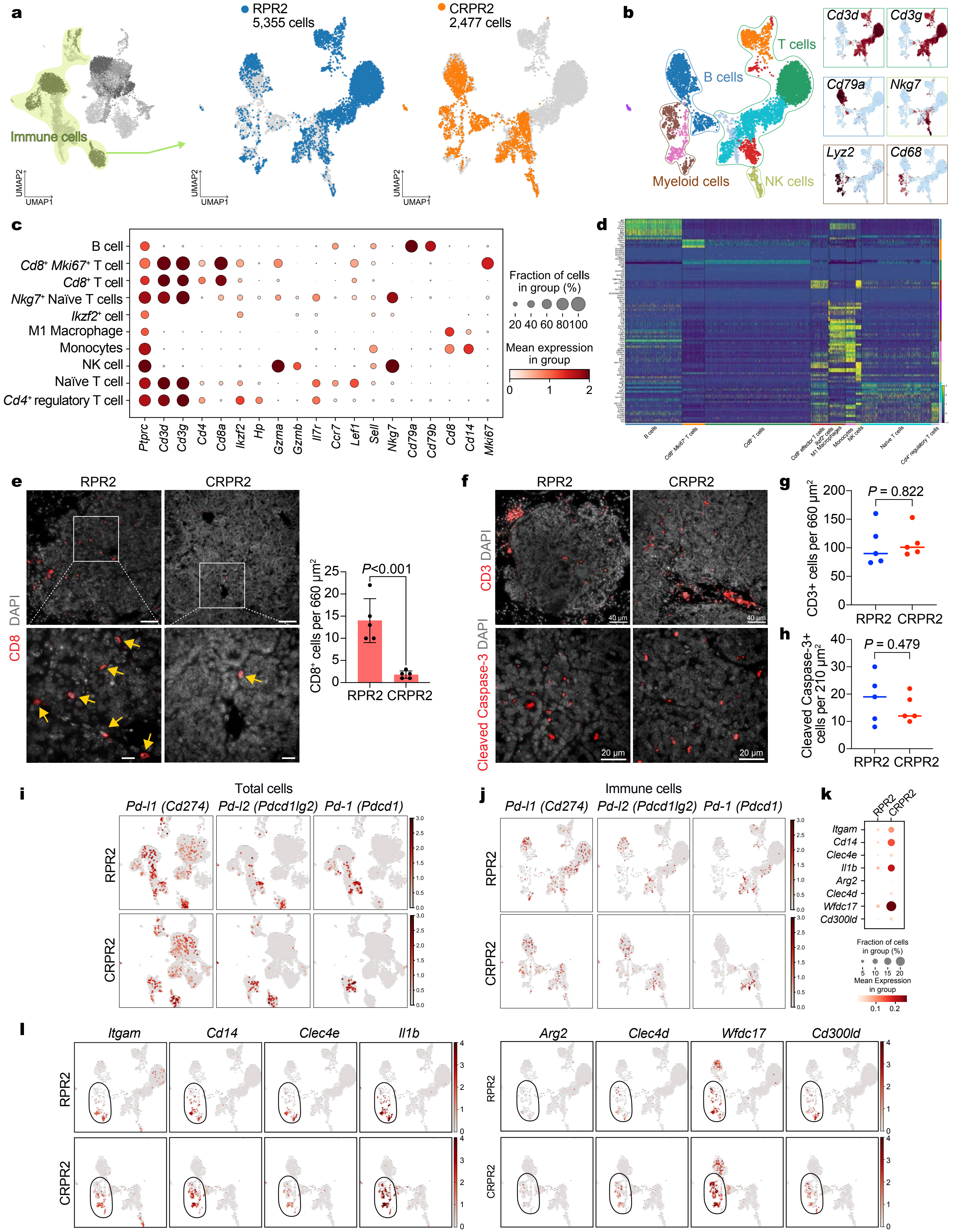

**Supplementary Figure 6**

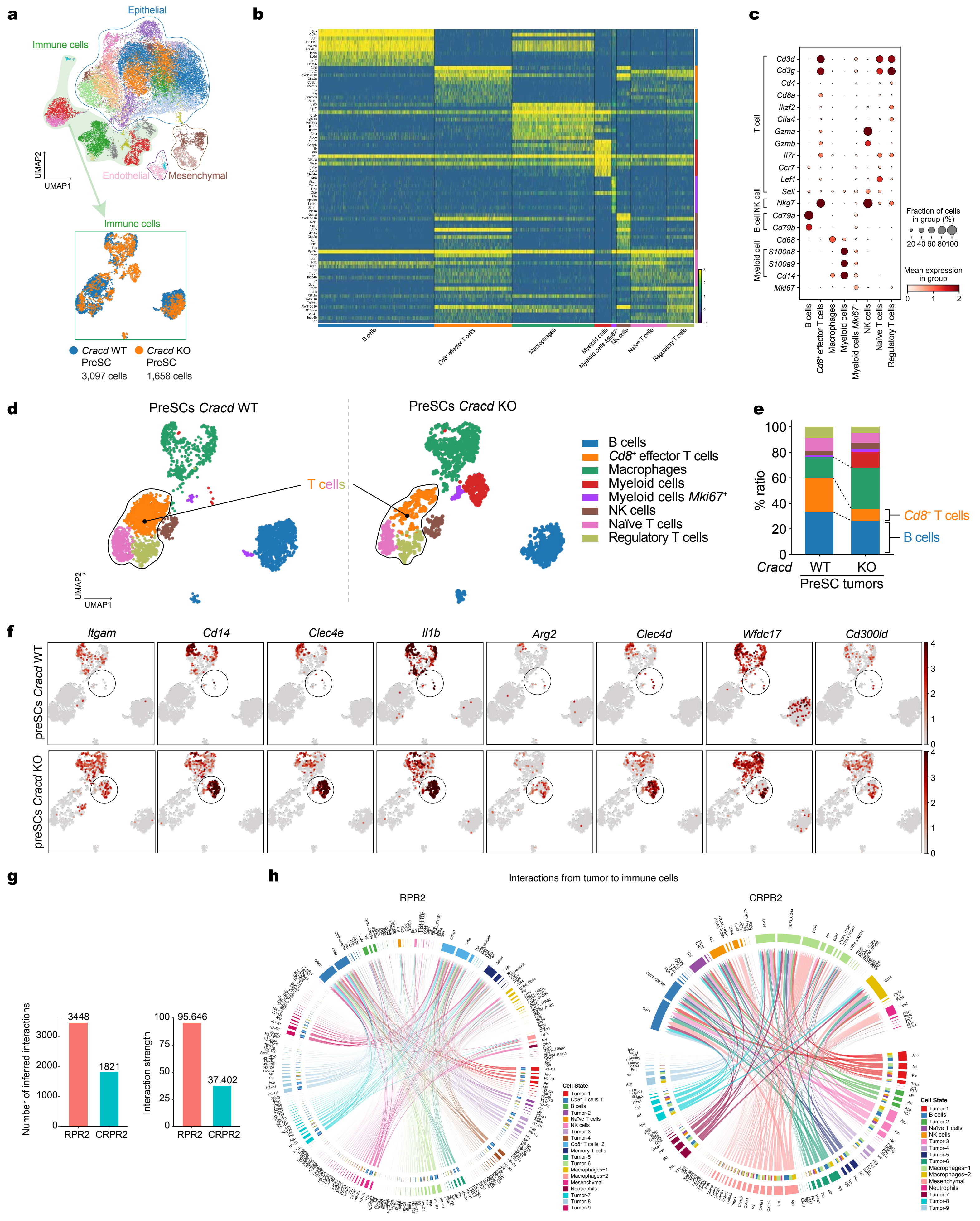

**Supplementary Figure 7**

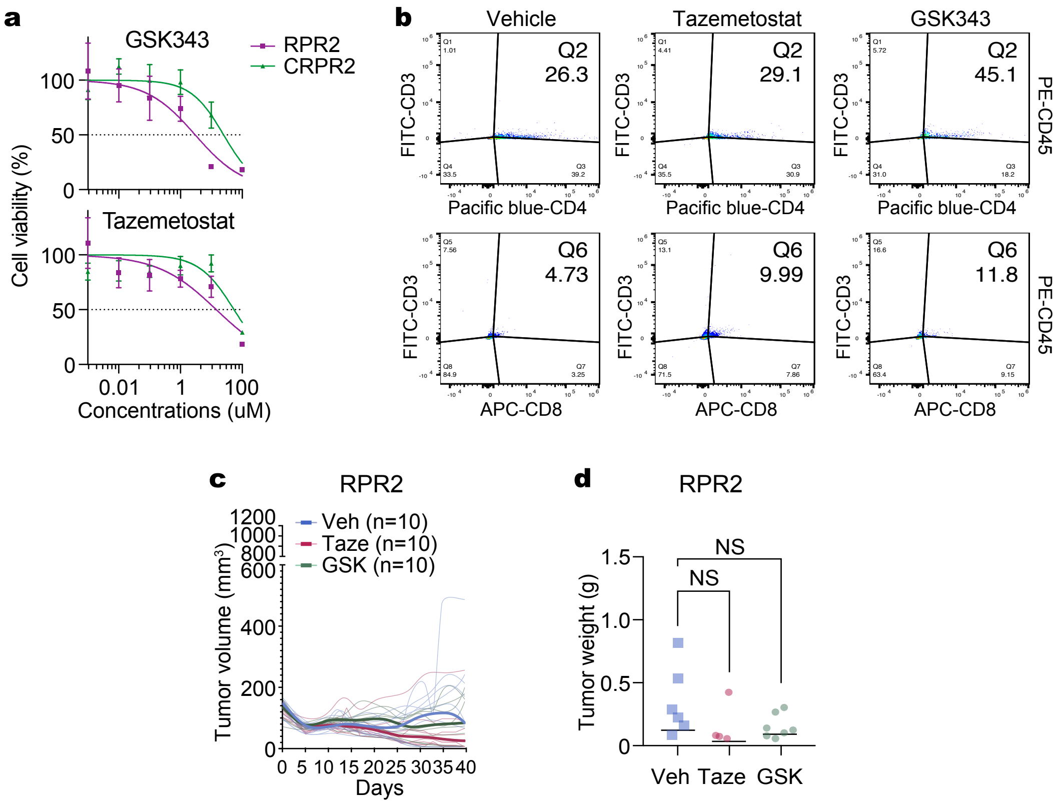

**Supplementary Figure 8**

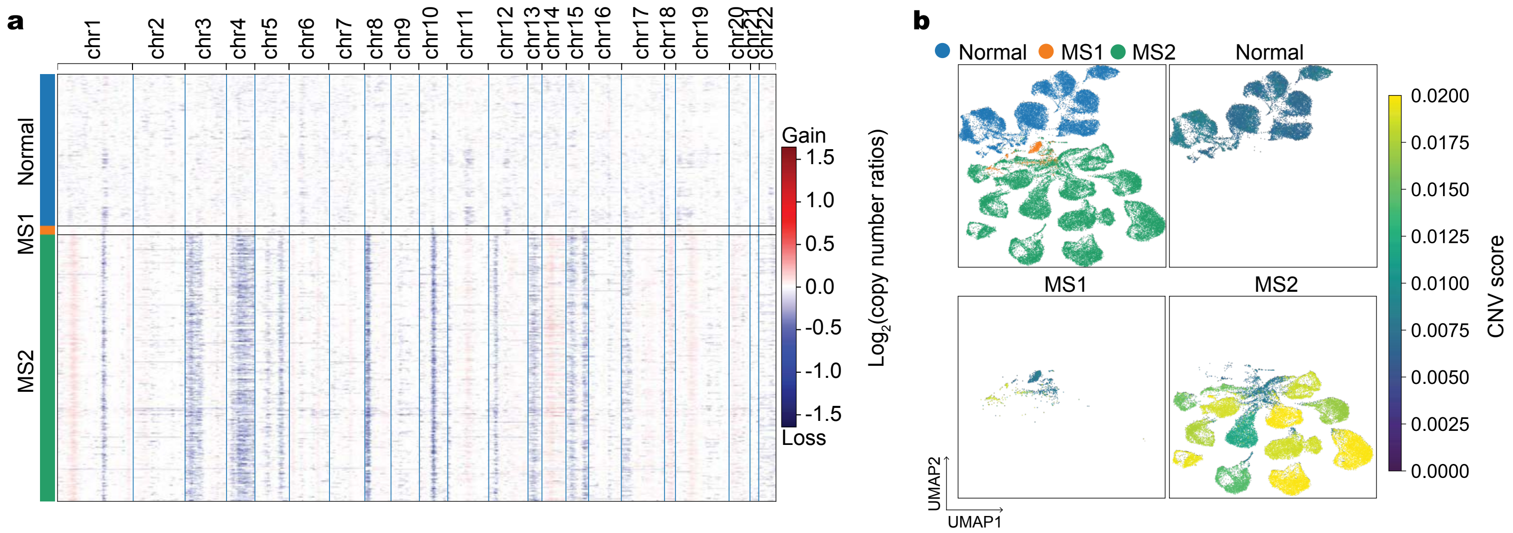
